# Integration in or Near Oncogenes Plays Only a Minor Role in Determining the in Vivo Distribution of HIV Integration Sites Before or During Suppressive Antiretroviral Therapy

**DOI:** 10.1101/2020.11.25.397653

**Authors:** John M. Coffin, Michael J. Bale, Daria Wells, Shuang Guo, Brian Luke, Jennifer M. Zerbato, Michele D. Sobolewskii, Twan Sia, Wei Shao, Xiaolin Wu, Frank Maldarelli, Mary F. Kearney, John W. Mellors, Stephen H. Hughes

## Abstract

HIV persists during antiretroviral therapy (ART) as integrated proviruses in cells descended from a small fraction of the CD4+ T cells infected prior to the initiation of ART. To better understand what controls HIV persistence and the distribution of integration sites (IS), we compared about 16,000 and 54,000 IS from individuals pre-ART and on ART, respectively, with approximately 385,000 IS from PBMC infected in vitro. The distribution of IS in vivo is quite similar to the distribution in PBMC, modified by selection against proviruses in expressed genes, by selection for proviruses integrated into one of 6 specific genes, and by clonal expansion. The clones in which a provirus integrated in an oncogene contributed to the survival of the clone comprise only a small fraction of the clones that persist in HIV-infected individuals on ART. Mechanisms that do not involve the provirus, or its location in the host genome, are more important in determining which clones expand and persist.

**Author Summary:** In HIV-infected individuals, a small fraction of the infected cells persist and divide. This reservoir persists on ART and can rekindle the infection if ART is discontinued. Because the number of possible sites of HIV DNA integration is very large, each infected cell, and all of its descendants, can be identified by the site where the provirus is integrated (IS). To understand the selective forces that determine the fates of infected cells in vivo, we compared the distribution of HIV IS in freshly-infected cells to cells from HIV-infected donors sampled both before and during ART. We found that, as has been previously reported, integration favors highly-expressed genes. However, over time the fraction of cells with proviruses integrated in highly-expressed genes decreases, implying that they grow less well. There are exceptions to this broad negative selection. When a provirus is integrated in a specific region in one of six genes, the proviruses affect the expression of the target gene, promoting growth and/or survival of the cell. Although this effect is striking, it is only a minor component of the forces that promote the growth and survival of the population of infected cells during ART.

## Introduction

Integration of viral DNA into the host cell genome is an essential step in the replication of HIV. In the course of untreated infection, the majority of infected cells dies within a few days [1-3], but a small fraction persists for many years of suppressive antiretroviral therapy (ART) and a small fraction of the persistent proviruses is the source of the virus that re-emerges when therapy is interrupted [4]. Understanding the mechanism(s) by which the infected cells persist is central to developing and evaluating strategies to cure HIV infections. Recent studies describing the distribution of integration sites (IS) in cells obtained from infected people have provided important clues about the persistence of infected cells (reviewed in [5, 6]).

The distribution of proviruses in HIV-infected individuals on long-term ART derives from their initial distribution in newly infected cells, modified by selection that promotes their survival, preferential loss, or clonal expansion. In vitro experiments show that the IS for HIV proviruses are widely distributed in the human genome; however, the distribution is far from random, favoring the bodies of highly-expressed genes located in gene-dense regions [7-9]. Retroviral integration exhibits a modest preference for specific bases around the site of integration [10, 11], and there is no discernable preference for the integration of the viral DNA in either orientation relative to the direction of transcription of the target gene [7, 12, 13].

There are good reasons to expect that HIV DNA integration follows the same rules in cells infected in culture and in a human host. In untreated chronic HIV infection, between one in 100 and one in 1000 CD4+ T cells are infected every day; almost all of the newly infected cells die. In untreated individuals, there are more than 10^8^ new integration events every day, a process that continues for many years. Only a small fraction of the infected cells survives for more than a few days following the initial infection. Most of the surviving cells (>98%) carry defective proviruses [14, 15]. However, despite the fact that the cells that carry intact infectious proviruses represent only a small fraction of the cells that survive, that still adds up to a large number of cells that carry infectious proviruses [15]. At least some of the infectious proviruses are in a poorly defined, but reversible, state of latency. These proviruses persist for decades of ART and can produce progeny viruses that can rekindle the infection when ART is interrupted [16, 17]. Collectively, the intact infectious proviruses constitute the viral reservoir.

The reservoir comprises proviruses in cells that were infected prior to the initiation of ART (and their descendants), and is not the result of persistent viral replication in blood or solid tissues [4, 18-20]. Studies of the distribution of IS in cells from infected individuals revealed that at least 40% of the long-lived, persistently infected cells were in a small number of clones. Clonally amplified cells can persist for at least 11 years [21-23]. There are clonally amplified CD4+ T cells that carry infectious proviruses, some of which express infectious viruses with potential to rekindle the infection when ART is discontinued [22, 24-27].

Although we now know that there is extensive clonal expansion of infected cells in vivo, important questions remain: 1) Integration in at least three genes (*MKL2, BACH2, STAT5B*) [21, 23, 28] can promote the survival and/or proliferation of HIV-infected cells. Are there other genes in which proviral DNA provides a selective advantage to the infected cell? 2) Are there other factors that help to reshape the initial distribution of IS in individuals on long term ART? 3) What is the relative importance of provirally-mediated expression of oncogenes and other factors (e.g., the immune response) in the clonal expansion and persistence of HIV-infected cells?

To address these questions, we prepared a large IS library from stimulated PBMCs infected in culture with HIV [29]. We then compared the distribution of the IS to the patterns of gene expression in these cells and to the distribution of IS obtained from cells from infected people both before ART and during suppressive ART.

## Results

### Study participants

IS data were obtained from the donors described in the studies listed in Table S1, and their characteristics can be found in the references. All studies had the approval of the relevant institutional IRB, and informed consent was obtained for all the samples collected, as described in the cited studies.

### Integration site datasets

The distribution of IS in people living with HIV on long-term ART can only be properly understood in the context of the distribution of IS at the time the cells were initially infected. Unfortunately, it has not been possible to prepare sufficiently large IS libraries from newly-infected people [18]. We therefore analyzed the distribution of HIV IS in PHA-stimulated CD8-depleted PBMC isolated from two HIV-negative individuals. The PBMC were infected in vitro with replication competent HIV-1_BAL_ (referred to hereafter as “PBMC”). We compared that distribution to the distribution of IS in cells obtained from infected individuals both pre-ART and on-ART using identical methods. The comparative analysis of the three datasets allowed us to assess the effects of various selective forces – both positive and negative – that modify the distribution of IS during suppressive antiretroviral ART.

Total DNA was extracted from the PBMC two days following infection, which should not allow time for selection for, or against, proviruses integrated in specific sites or specific genes [29]. We used linker-mediated PCR [30] as previously described [21, 31] to identify IS. When we compared the global distribution of the IS from the PBMC obtained from the two individuals, the correlation coefficient was 0.98 and, for the rest of the analyses, we combined the data from the two sets of IS. The combined dataset from the in vitro-infected PBMC contained 394,975 independent IS. It will be referred to as the “PBMC dataset” for the remainder of the pape

By combining in vivo IS from several studies [18, 21, 32, 33] (Table S1), we were able to assemble datasets comprising a total of 15,780 IS from pre-ART samples from 30 HIV-infected donors (hereafter referred to as “donors”) and 53,945 IS from on-ART samples from 39 donors (Table 1, S1). The introduction of a shearing step prior to linker ligation and PCR [30] creates different break points in flanking host DNA, making it possible to distinguish identical IS that arose by clonal expansion of a single infected cell [21, 30, 31]. After collapsing the IS with multiple breakpoints to a unique site, the two libraries contained 13,324 unique IS pre-ART and 33,336 unique IS on-ART (Table 1).

**Table 1.**
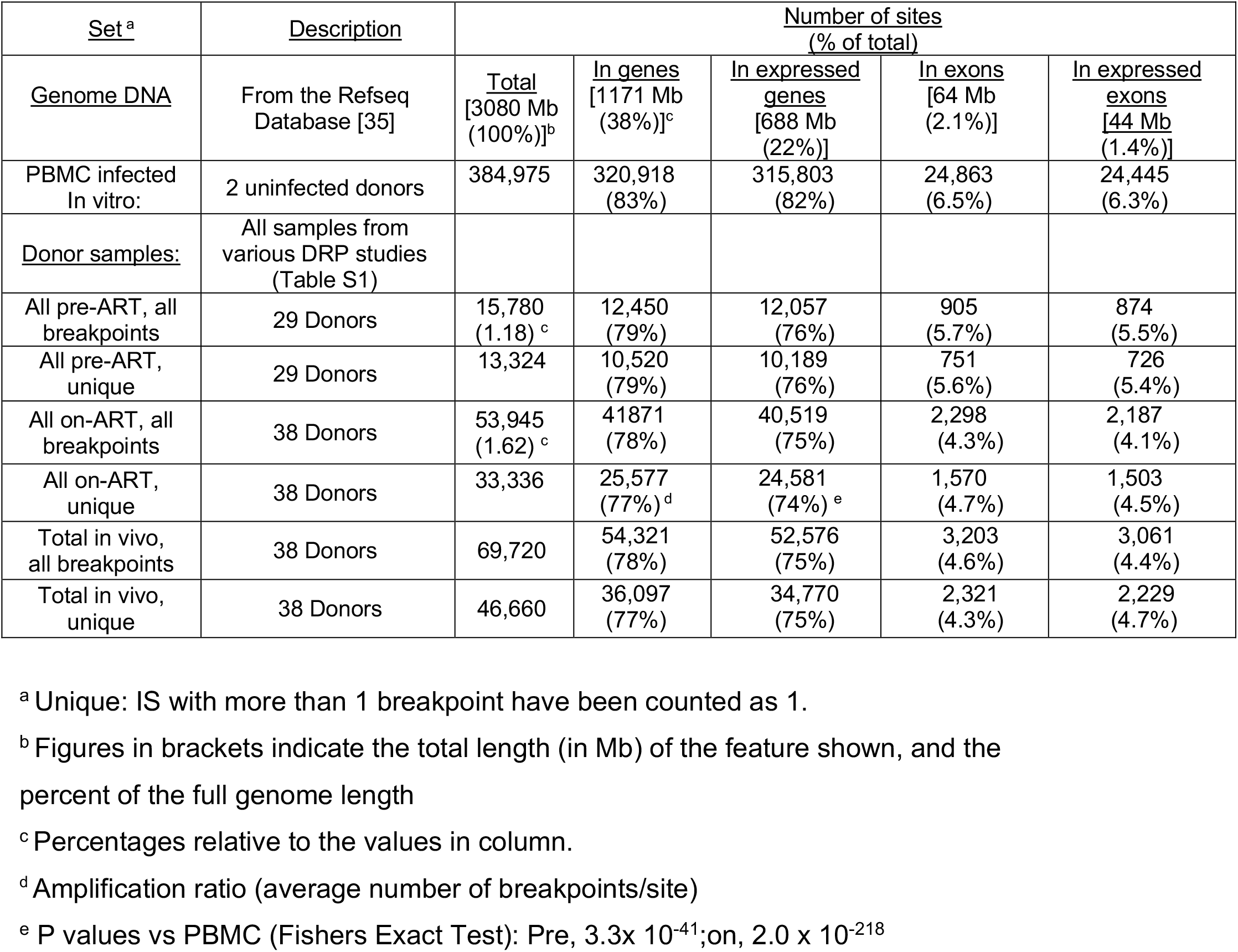
Integration datasets used in this study.

### In vivo selection against proviruses integrated in highly-expressed genes

HIV preferentially integrates a DNA copy of its genome into highly-expressed genes in gene-rich regions [8, 9, 34]. However, the initial analyses were based on small datasets (<1 IS/gene), used methodology (restriction enzyme cleavage) which has the potential to introduce bias based on local base composition differences, and used cell lines rather than primary cells. To confirm the prior results and extend them to relevant target cells, we mapped the IS in the PBMC dataset into genes from the entire RefSeq gene database [35], slightly modified to remove gene overlaps, yielding a set of 20,207 genes. The overlaps were removed to prevent mapping single IS to two genes. To compare the sites of integration to the expression of the host genes, we performed an RNA-seq analysis using the same PBMC that were used to generate the PBMC IS dataset. Table 1 summarizes the results of this analysis. As expected, most of the IS (83%) were in genes, which comprise only 38% of the human genome, and most of the IS (82%) were in expressed genes, which comprise about 22% of the genome. Thus, only 18% of the HIV IS were found in the 78% of the genome that is not in expressed genes. A similar preference was evident in the pre-ART data in which 79% of the IS were in genes and and 76% were in expressed genes. In the on-ART dataset 77% of the unique IS were in genes and 74% of the pre-ART sites were in expressed genes. Although only 2.1% of the genome is in exons, exonic IS comprised 6.5% of the IS in PBMC and somewhat less (5.6% and 4.7%) of the unique pre- and on-ART samples, respectively.

The differences between the percentages of the IS in genes in the in vitro PBMC and in the on-ART or pre-ART datasets, although not large (4% and 7%), were highly significant (Table 1). There is an even larger difference in expressed genes (7% and 9%). We compared the relationship of IS to gene expression in the three datasets using our non-overlapping gene set (available in Supplemental File 1). All RefSeq genes were divided into 100 bins based on the RNA-seq data (as transcripts per million reads, TPM) obtained from the HIV-infected PBMC (Figure 1, green triangles) and also divided into 4 bins, again based on TPM. The mean integration site density (unique IS/Mb, normalized to the mean for each dataset) for the 3 IS datasets when the IS are dividied into the 4 bins, is presented in Figure 1A. The same data, divided into 100-bins are shown in Figure S1. As expected [8, 9, 34], the results showed a strong correlation between integration site density and gene expression for all three datasets. There was a small, but highly significant, difference in the integration site density in the 3 datasets for the genes with the highest TPM. When the IS data were divided into 4 bins, the fraction of IS that were in the quartile of the most highly-expressed genes was about 10% lower in the pre-ART dataset than in the PBMC dataset, and about 18% lower in the on-ART dataset, compared to the PBMC dataset (p =3.3 x 10^-41^). A smaller effect was seen in the second quartile, based on gene expression (p =2.0 x 10^-218^). No similar effect was seen in the genes in the bins with two lowest levels of expression. This result implies the presence of selection, in vivo, against cells containing proviruses integrated in the most highly-expressed genes, and shows that the selection operated both pre-ART and on ART.

**Figure 1.**
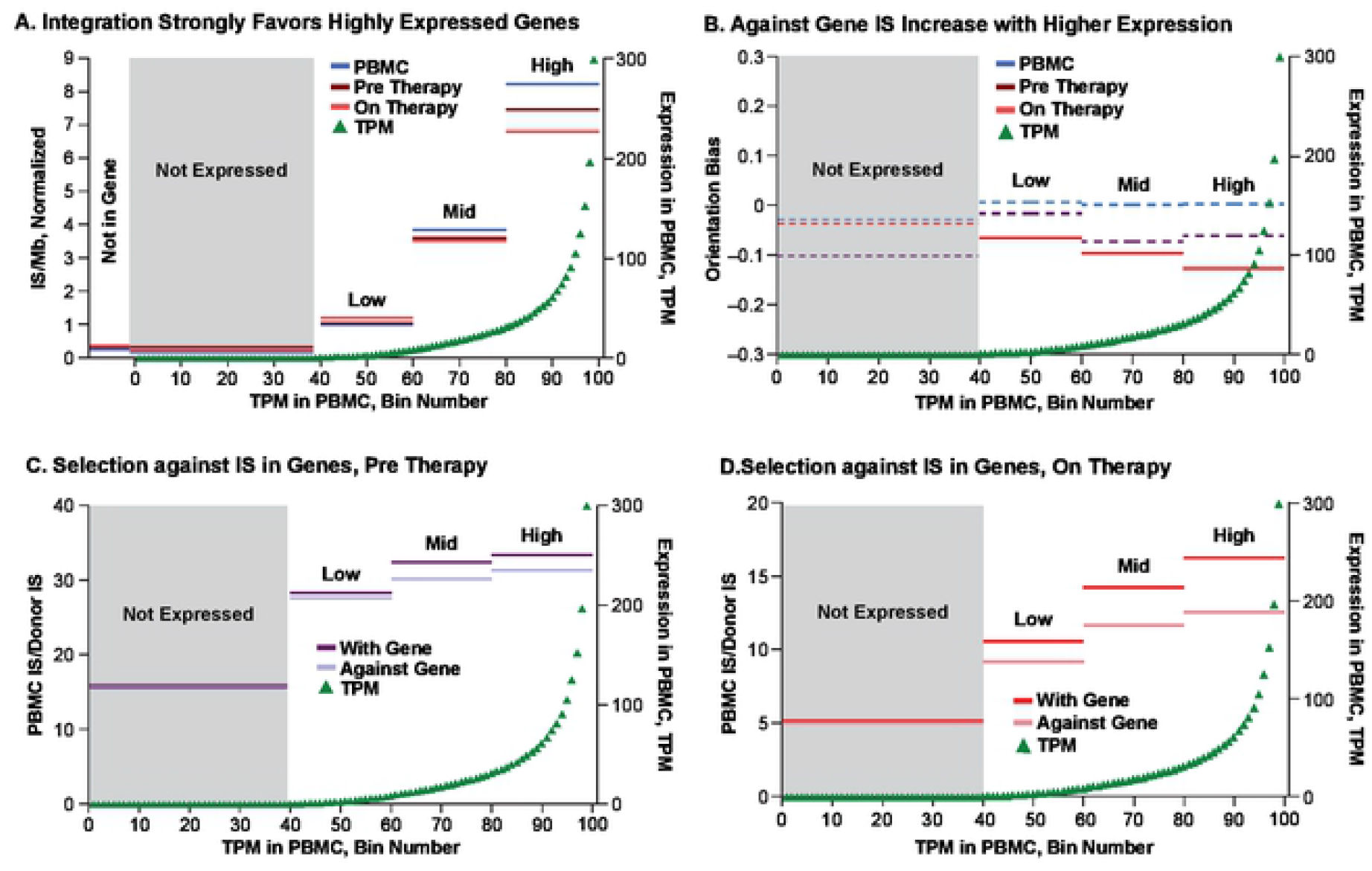
Gene expression and integration. The complete set of ca 22,000 RefSeq genes was slightly modified to remove overlaps (see Materials and Methods). The non-overlapping genes were divided into either 100 bins or 4 bins, based on the RNA-seq analysis (TPM) of the in vitro infected PBMC. The 100-bin RNA-seq data are shown in green triangles in all panels. The combined IS data are shown for the genes in each of the 4 bins in all panels for PBMC (blue), pre-ART donors (plum) and on-ART donors (red). Darker colors in C and D indicate proviruses oriented in the same direction as the host gene; lighter colors are in the opposite orientation. **A**. Total IS density (sites/Mb) in each bin, normalized to the average for the whole genome (125 sites/Mb for PBMC, 4.28 sites/Mb pre-ART, and 10.7 sites/Mb on-ART). **B**. The orientation of the proviruses relative to the host gene was calculated for each bin as (proviruses with gene-proviruses against gene)/(total proviruses). Dashed lines indicate p values (binomial) >0.05. **C** and **D**. Ratios of proviruses per bin for the pre-ART (**C**) or on-ART (**D**) samples. Note that the higher the ratio, the smaller the number of IS in the donor samples relative to the in vitro infected PBMC samples. Figure S1 shows the same results with the 100-bin IS data included.

We and others previously reported a preference for proviruses integrated in an orientation opposite to the host gene in HIV-infected individuals and SIV-infected macaques on-ART (5% and 9%, respectively) that is not seen in vitro [21, 23, 29]. When we separated the IS in genes according to their orientation relative to the host gene, there was, as expected, no significant bias for the orientation of the proviruses, relative to the host gene, in the PBMC dataset (Figure 1B). However, enrichment for proviruses orientated opposite to the on-ART dataset relative to the PBMC dataset was evident (Table S2, p=1.7 x 10^-63^). We also observed a smaller, but significant, enrichnment for proviruses in the opposite orientation to the gene in the pre-ART data (Table S2, p=0.002).

To understand the nature of the in vivo selection, we separated the IS data, based on whether the provirus was oriented with the gene, or against the gene, and the plotted IS data relative to the level of expression of the host gene, dividing the genes into four bins based on their levels of RNA expression. There was no significant selection against proviruses that were in the same ortientaion as the host gene in genes that were not expressed in either the pre-ART or the on-ART dataset (Figure 1B). In expressed genes, there was a significant selection against proviruses oriented with the gene in the on-ART dataset [p values ranging from about 0.013 to 5 x 10^-47^ (Table S2)]; the strongest selection was seen in the highly-expressed genes. A similar pattern was seen with the pre-ART data, although the effects were smaller and the effect for the genes that were expressed at a low level was not significant. While the difference in IS distribution between the PBMC and the in vivo samples could have reflected differences in initial integration preferences between the two conditions, the differences between pre- and on-ART samples can only be explained by selective forces that affect the survival or proliferation of the infected cells. The on-ART data (Figure 1D) also showed that there is weaker selection against cells with proviruses integrated in highly-expressed genes in the opposite orientation to the gene (p=2.3 x 10^-48^); again the strength of selection decreased with decreasing levels of gene expression. A similar, but much weaker, selection was seen in the pre-ART dataset (p=0.011).

### Selection involving proviruses integrated in individual genes

To examine the contribution of proviruses in genes that are favored for HIV integration (hot spots) we ranked genes in both the PBMC and the on-ART datasets by IS density, as unique sites per kb (Table 2). For the most part, the IS site density in individual genes showed that the in vitro and in vivo data were in good agreement (Table 2, Figure 2). We and others previously reported that proviruses integrated in certain introns of three genes (*BACH2, MKL2, STAT5B*) can confer a selective advantage for the clonal expansion and/or survival of infected cells [21, 23, 28]. The in vivo enrichment of IS in these genes is due to post-integration selection, an effect that has sometimes been misinterpreted in the literature as reflecting preferential “hot spots” for integration [36, 37].

**Table 2.**
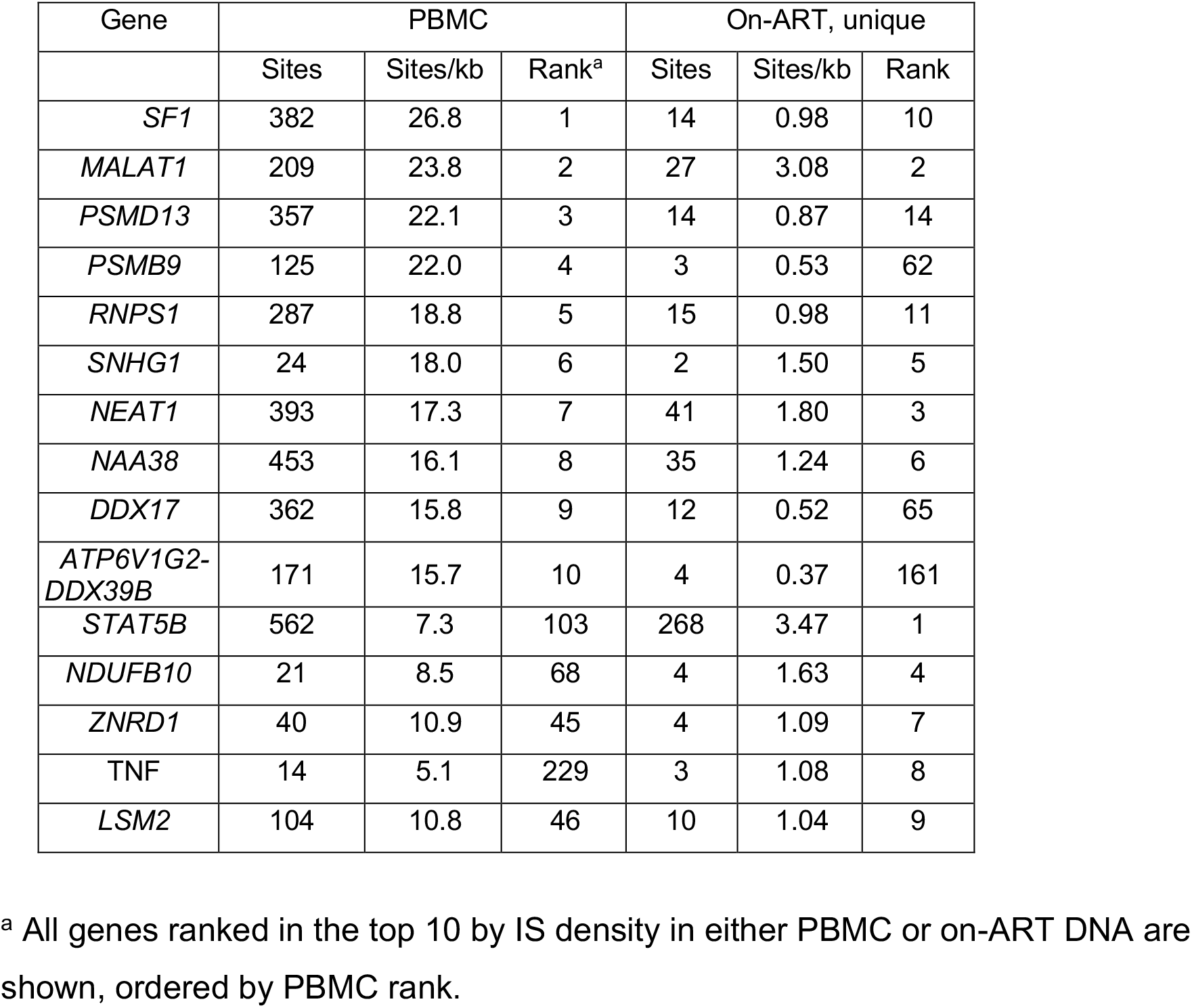
Genes favored as targets for integration in PBMC infected in vitro and the on-ART samples.

| Gene | PBMC |  |  | On-ART, unique |  |  |
| --- | --- | --- | --- | --- | --- | --- |
|  | Sites | Sites/kb | Rank<sup>a</sup> | Sites | Sites/kb | Rank |
| *SF1* | 382 | 26.8 | 1 | 14 | 0.98 | 10 |
| *MALAT1* | 209 | 23.8 | 2 | 27 | 3.08 | 2 |
| *PSMD13* | 357 | 22.1 | 3 | 14 | 0.87 | 14 |
| *PSMB9* | 125 | 22.0 | 4 | 3 | 0.53 | 62 |
| *RNPS1* | 287 | 18.8 | 5 | 15 | 0.98 | 11 |
| *SNHG1* | 24 | 18.0 | 6 | 2 | 1.50 | 5 |
| *NEAT1* | 393 | 17.3 | 7 | 41 | 1.80 | 3 |
| *NAA38* | 453 | 16.1 | 8 | 35 | 1.24 | 6 |
| *DDX17* | 362 | 15.8 | 9 | 12 | 0.52 | 65 |
| *ATP6V1G2-DDX39B* | 171 | 15.7 | 10 | 4 | 0.37 | 161 |
| *STAT5B* | 562 | 7.3 | 103 | 268 | 3.47 | 1 |
| *NDUFB10* | 21 | 8.5 | 68 | 4 | 1.63 | 4 |
| *ZNRD1* | 40 | 10.9 | 45 | 4 | 1.09 | 7 |
| *TNF* | 14 | 5.1 | 229 | 3 | 1.08 | 8 |
| *LSM2* | 104 | 10.8 | 46 | 10 | 1.04 | 9 |

<sup>a</sup> All genes ranked in the top 10 by IS density in either PBMC or on-ART DNA are shown, ordered by PBMC rank.

**Figure 2.**
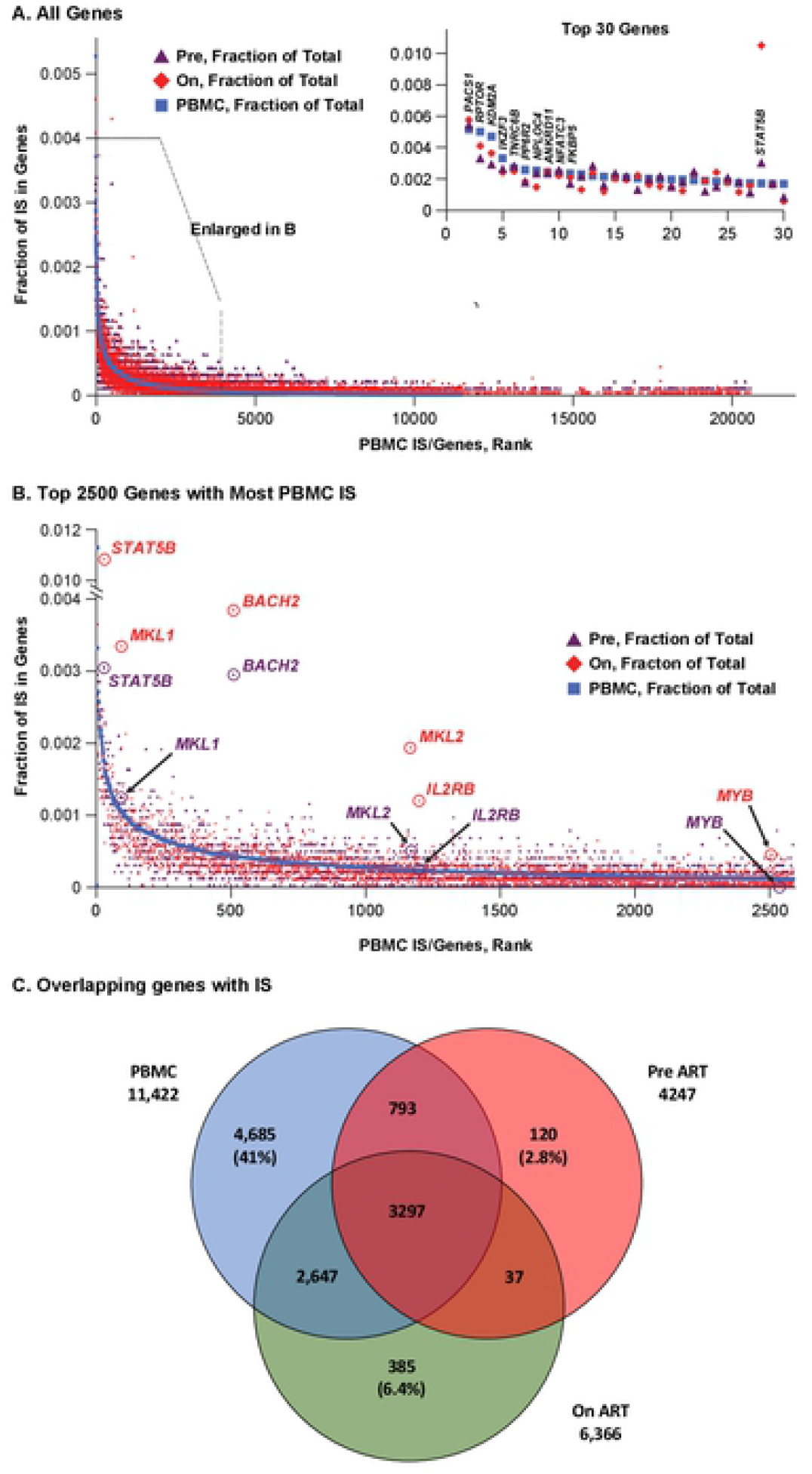
Comparative ranking of genes by integration frequency. The non-overlapping RefSeq genes were ranked according to the number of unique IS in the PBMC in vitro dataset (blue squares), along with the number of sites in the same genes in the pre-ART (plum triangles) and on-ART (red diamonds) datasets. **A**. All 20,000 genes in the dataset are shown; about half have one or more IS in the in vitro PBMC dataset. Points with no IS have been removed for clarity, and the genes with no IS in PBMC are assigned a rank at random. The inset shows an expanded view of the 30 genes with the most IS. The top 10 are labeled, as is *STAT5B* (number 28). **B.** Expanded view of the 2500 genes (indicated by the box in **A**) with the most IS. The 6 genes in which an integrated provirus directly contributes to persistence and/or clonal expansion of the infected cell are circled and labeled. Some pre-ART points are indicated with arrows for clarity. **C.** Venn diagram showing the genes with at least one IS in the three datasets, and how those genes are shared among the datasets. Numbers indicate the number of genes in each category.

To ask if there are other genes in which an HIV provirus could provide a selective advantage to the host cell in vivo, we compared, for each gene, the fraction of IS in the in vitro PBMC dataset to the fraction in the pre-ART and on-ART in vivo datasets (Figure 2). The two panels show the same data using two different scales. Panel A shows the data for all of the genes. The inset shows the top 30 genes, showing that, for most genes, there is a strong correlation between the IS distribution in vitro and in vivo. There is one obvious exception to this correlation in the top 30 genes, *STAT5B*, a gene in which HIV proviruses are known to confer a selective advantage [28]. Panel B shows the top 2600 genes by IS frequency in PBMC. Although there is some scatter in the data, the in vivo IS data closely mirror the PBMC data.

Panel C is a Venn diagram that shows the extensive overlap between the genes that are favored for integration in the 3 datasets. Of 4,247 genes with at least one IS in the pre-ART dataset, 4090 (96%) also had at least one IS in the PBMC dataset (there are11,492 genes IS with at least one IS in PBMC). Similarly, of the 6366 genes in the on-ART dataset, only 422 (6.6%) had no IS in either of the other two sets, and 5099 (93%) of the genes with at least one IS in the on-ART dataset had at least one IS in PBMC. These comparisons validate the use of the in vitro PBMC data as a surrogate for the initial distribution of IS in vivo and show that the strongest determinant of the overall distribution of HIV proviruses in infected individuals, both pre-ART and on-ART, is the initial distribution at the time the cells (or their ancestors) were infected.

Despite the overall selection against proviruses in expressed genes, there were a few genes, circled in Figure 2B, in which there was a higher fraction of proviruses in vivo than in vitro, implying that there is an in vivo selection for cells with HIV proviruses in these genes. As will be discussed below, proviruses in these genes show other evidence for a positive in vivo selective effect in the host cells–clustering of IS within the gene and a strong bias toward selection for proviruses in the same orientation as the gene (Table 3).

**Table 3.** Genes in which integrated provirus can be selected in vivo.

| Gene name | Unique IS in genes on-ART | IS in PBMC | PBMC/on-ART Ratio | Enrichment probability (Fishers) | Provirus orientation: with gene/against | Orientation probability (Fishers) | Selected IS relative to protein coding introns |
| --- | --- | --- | --- | --- | --- | --- | --- |
| All genes | 25,731 | 326,033 | 12.67 | | 11,476/14,255 | $7.0 \times 10^{-60}$ | |
| *STAT5B* | 268 | 562 | 2.10 | $1.1 \times 10^{-108}$ | 197/71 | $6.6 \times 10^{-12}$ | Upstream |
| *BACH2* | 98 | 132 | 1.34 | $1.2 \times 10^{-43}$ | 71/20 | $2.4 \times 10^{-5}$ | Upstream |
| *MKL2* | 49 | 69 | 1.41 | $2.5 \times 10^{-24}$ | 40/6 | $2.8 \times 10^{-5}$ | In between |
| *MKL1* | 85 | 331 | 3.89 | $3.3 \times 10^{-10}$ | 53/30 | 0.02 | In between |
| *IL2RB* | 30 | 68 | 2.27 | $3.3 \times 10^{-10}$ | 17/9 | 0.041 | Upstream |
| *MYB* | 11 | 31 | 3.10 | $2.9 \times 10^{-4}$ | 10/0 | 0.0085 | In between |

### Mapping and comparing the distribution of integration sites in vivo and in vitro

To help compare the distribution of IS in different datasets, we developed an Excel-based tool which can be used to view different sized regions of the host genome ranging from a selected portion of a gene to a whole chromosome. The tool, available as a Supplemental File, makes it possible to compare the IS site distribution and gene expression levels for up to 5 IS datasets at a time (Figure S2). The Excel workbook, including the visualization tool, gene list, and the RNA-seq integration data used in this study is available in the supplemental material. The tool divides any chosen region of the genome into 250 bins. The number of IS per bin is shown as vertical bars; red indicates IS in the same orientation as the numbering of the chromosome, blue, the reverse orientation. The levels of RNA (TPM) in HIV-infected PBMC are shown as grey boxes. The locations of genes are indicated, as is their transcriptional orientation.

Figure 3 shows a comparison of the distribution of IS in the on-ART dataset and the in vitro PBMC dataset. We chose to show the data for chromosome 16 because it contains *MKL2*, one of the genes in which an HIV provirus can provide positive selection for the host cell in vivo (see Figure 2B) [21]. The IS data are shown at several scales, from the whole chromosome (bin size ca 355,000 bp, Figure 3A) down to a fraction of a gene (bin size 150 bp, see Figure 4B). At the three largest scales, the pattern of IS is strikingly similar in the PBMC and on-ART datasets, and closely matches the distribution of the expressed genes.

**Figure 3.**
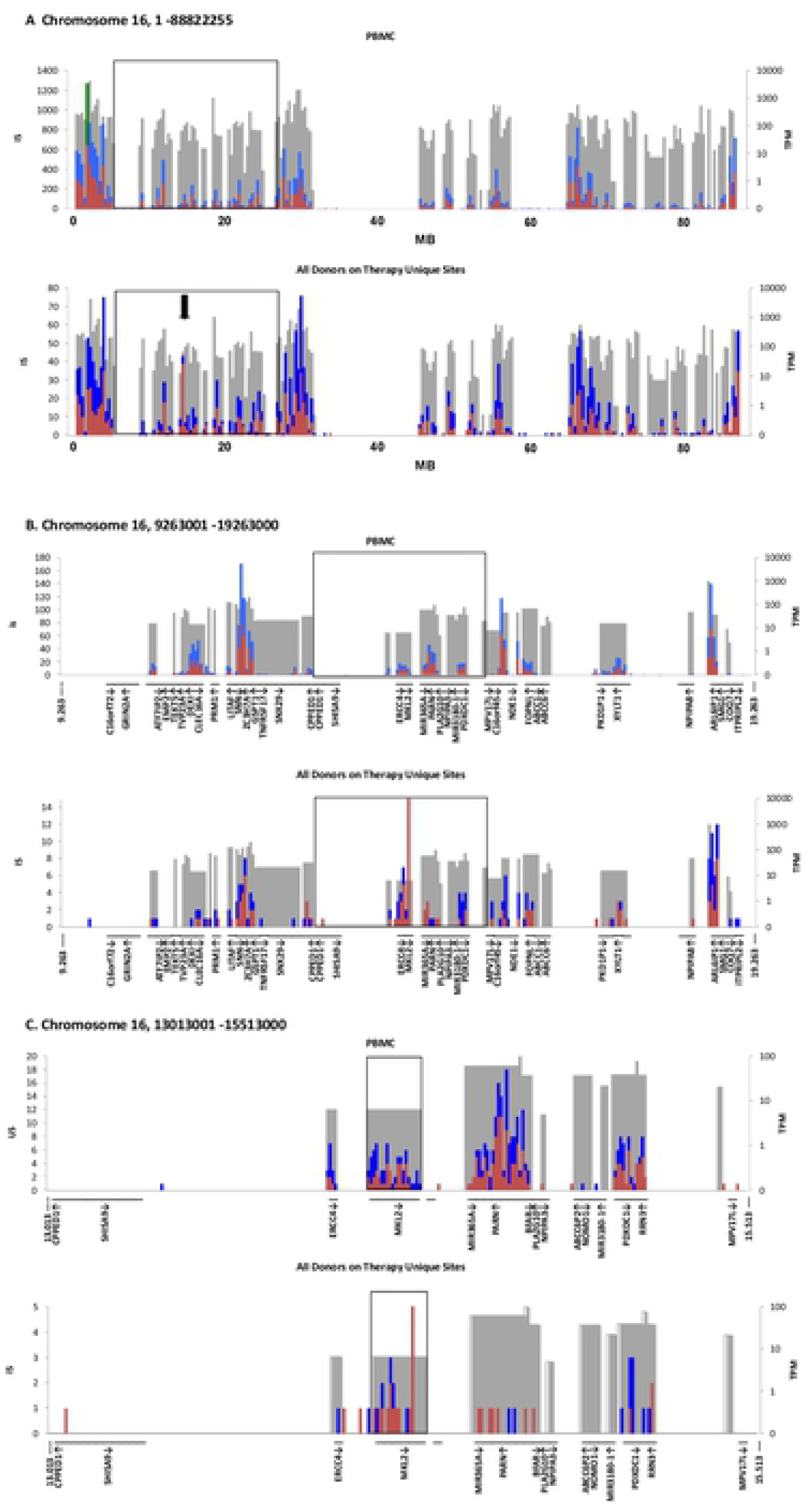
Analysis of IS in MKL2. Using the Excel-based application described in Figure S2, a selected region of the genome of any size is divided into 250 bins. The IS in each bin are tabulated and the number of IS in the bin is shown as a bar, with the orientation of the provirus relative to the numbering of the chromosome indicated by color (red for the same as the chromosome and blue for the opposite). The grey boxes indicate the location and relative RNA level of the genes in each bin. RefSeq genes (from the hg19 sequence database) are shown at the bottom of each plot, with arrows indicating their orientation relative to the numbering of the chromosome. The figure shows the distribution of genes and IS on chromosome 16 at 3 different scales. In all panels, the top image shows the distribution of IS from PBMC infected in vitro; the bottom, the distribution of the unique IS data from on-ART donors. **A.** Entire chromosome 16 (ca 353,000 bp/bin); the arrow shows the region in which IS are enriched in the on-ART samples. The box shows the region enlarged in panel B. **B.** A 10Mb region of chromosome 16, (40 kb/bin) centered on MKL2. Again, the box indicates the region expanded in panel C. **C.** A 2 Mb region of chromosome 16 (8 kb/bin) centered on *MKL2*. The boxed region is expanded in Figure 4A.

**Figure 4.**
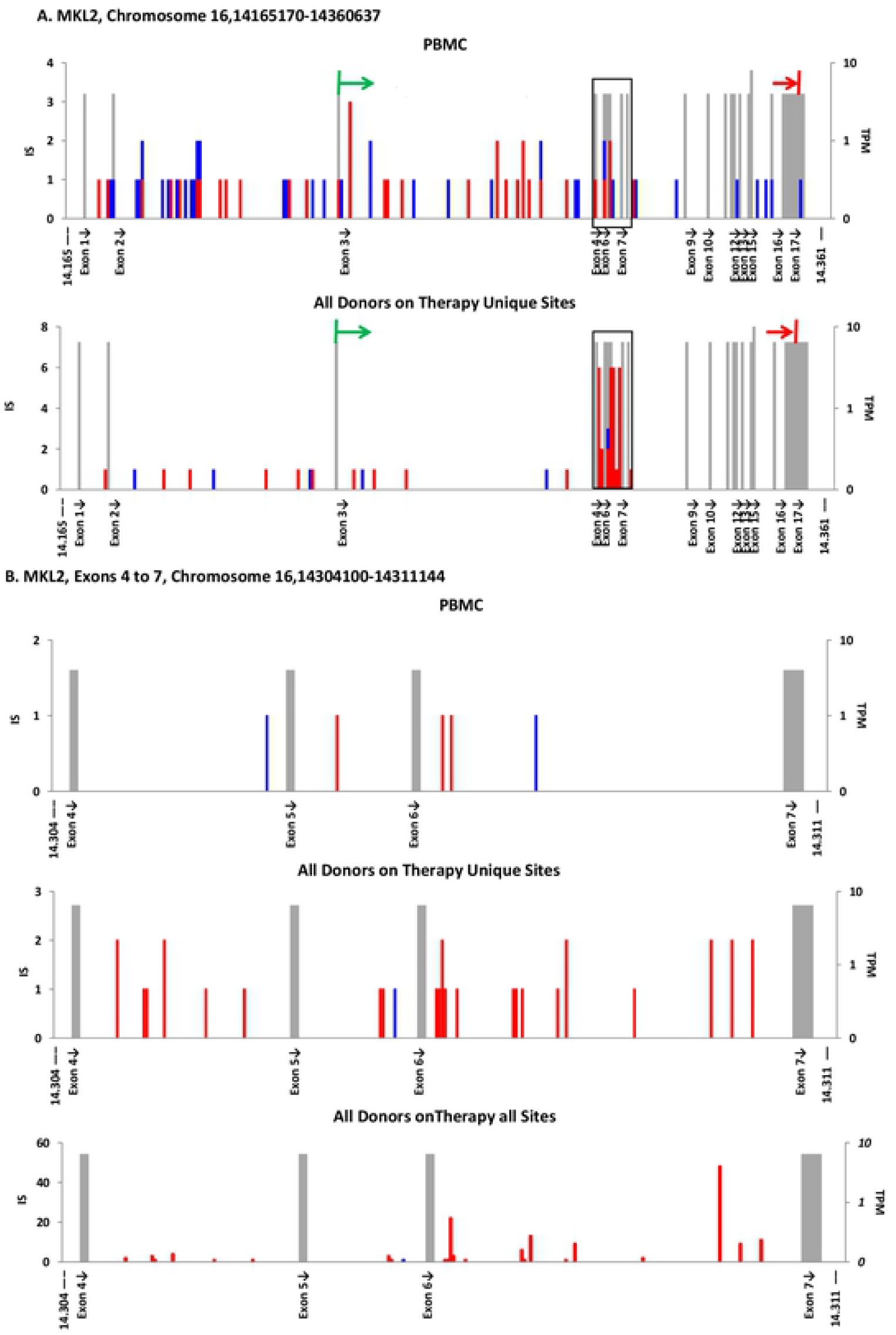
IS in *MKL2*. **A.** The complete *MKL2* gene (308 bp/bin). The gray bars show the position of exons, with their height indicating the expression level of the gene in TPM, as in Figure 3. The protein coding region is indicated by the colored arrows above the maps. The boxed region is enlarged in panel B. **B.** Cluster of *MKL2* IS on-ART in introns 4-6 (50bp/bin). Top, in vitro PBMC IS; middle, onART unique sites; bottom on-ART, all sites.

### Cclustered IS in specific genes

The one exception to the strong similarity in the distribution of the proviruses in chromosome 16 in the PBMC and on-ART datasets is in the *MKL2* gene (also known as *MTRFB*), marked by an arrow in the lower panel of Figure 3A. This difference maps to the *MKL2* gene (Figure 3B,C). A map of the *MKL2* gene, showing the location of exons in addition to the distribution of the IS, is shown in Figure 4A. As Figure 3 shows, this gene is not favored for integration in PBMC. However, in the on-ART dataset, proviruses are found in 30 different sites within a 7 kb region, encompassing introns 4-6 of the *MKL2* gene (Figure 4B). Most of these proviruses were in clonally amplified cells (Figure 4B, compare the bottom 2 panels). The probabilities that the cluster would exist by chance or that almost all of the proviruses in the cluster would be in the same orientation are both extremely small (2.5 x 10^-24^ and 3 x 10^-5^, respectively). These data support our previous conclusion that this cluster of IS is the result of selection for cells in which a provirus integrated in a small intronic region of *MKL2* altered the normal expression and/or structure of the *MKL2* protein. Misexpression of the *MKL2* protein would, in turn, provide a selective advantage to cells carrying these proviruses.

In addition to *MKL2*, HIV proviruses integrated in two other genes, *BACH2* and *STAT5B* have been previously shown to contribute to the growth and persistence of the infected cells in indivduals on ART [21, 23, 28]. Proviruses in these genes share characteristics with the *MKL2* proviruses: enrichment for the proviruses in vivo relative to the frequency seen in PBMC infected in vitro, clustering of the proviruses in one or a few neighboring introns, and the orientation of the proviruses is the same as the host gene (Figure 5 and Table 3). The current dataset provides strong statistical support for selection of proviruses integrated in *BACH2* and *STAT5B* on-ART (Table 3), and there is weaker evidence of selection for proviruses in some of these genes pre-ART (Figure 2).

**Figure 5.**
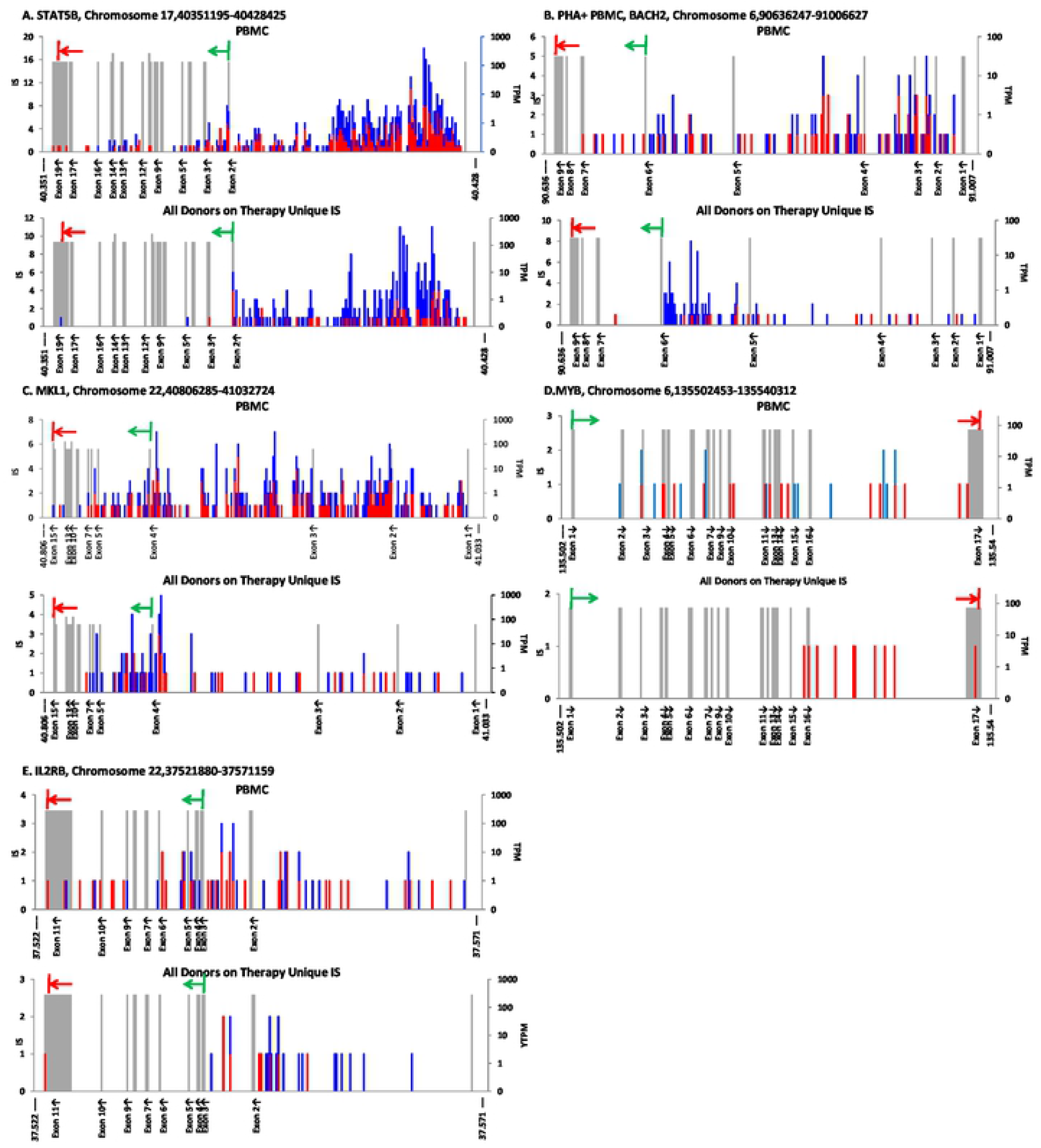
HIV IS in other genes in which proviruses can give the host cell a selective advantage. Maps were generated for each of the genes in Table 3 as described in Figure 4. For each pair of panels, the map at the top shows the IS distribution for the PBMC infected in vitro, and the on-ART distribution is at the bottom **A.** *STAT5B* (309 bp/bin). **B.** *BACH2* (1.48 kb/bin). **C.***MKL1* (906 bp/b**D.** *MYB* (151 bp/bin). **E.** *IL2RB* (197 bp/bin).

It was possible that there are additional genes in which HIV proviruses could provide a selective advantage for the host cell. We analyzed the total on-ART dataset for the three hallmarks of selection for cells with proviruses in specific genes: 1) enrichment of the IS in the on-ART dataset; 2) enrichment in the on-ART dataset for proviruses in the same orientation as the gene; and 3) clustering of the IS in one or a few nearby introns (Table 3).

As expected, when all 5983 genes with IS in both the on-ART and the PBMC datasets were analyzed and ranked by the combined p value for the relative number of IS and for orientation bias, the three top ranking genes were *MKL2, STAT5B*, and *BACH2*. Three more genes were identified using the new on-ART dataset: *MKL1* (also known as *MRFTA*), a paralogue of *MKL2* [38]; *IL2RB*, encoding the β chain of the IL-2 receptor (https://www.ncbi.nlm.nih.gov/gene/3560); and *MYB*, a known protooncogene originally identified in an avian retrovirus [39]. All six genes have been linked directly to cell growth or survival (see Discussion). In all of these genes, the proviruses in the on-ART dataset displayed a strong bias for integration in the same orientation as the gene and the number of IS was significantly greater than what would have been predicted based on the PBMC dataset (Figure 5, Table 3). In each of the six genes, the IS in the on-ART datasets showed evidence of clustering in specific introns. In the current on-ART dataset, these six genes were the only ones for which there was significant evidence for positive selection of cells with an integrated HIV provirus in vivo. Taken together, the 541 independent IS in the six genes in the on-ART data comprised about 1.6% of the unique IS in the dataset. Because our on-ART dataset is large, we were able to identify genes for which the number of IS represented a very small fraction of the total IS (*MYB* is 10/33,000 unique IS, or 0.00030). There may be additional genes that we have not identified in which proviral integration could lead to positive selection; however, the fraction of the total IS in such genes must be extremely small (see Discussion).

In the analysis of the on-ART dataset, we identified a number of genes for which there appeared to be selection for proviruses oriented opposite to the host gene. In 6 out of 8 cases, these genes had fewer IS in the on-ART dataset than would be expected based on the PBMC dataset (Table S3), consistent with stronger than average selection against cells with proviruses integrated in the same orientation as the gene. Unlike the genes in which there was evidence of selection for a provirus in the same orientation as the gene, in the six genes for which there appeared to be negative selection, the proviruses were not obviously clustered; the overall distribution of IS within the genes appeared to be similar to that seen in the PBMC dataset (Figure S3). One of the 6 genes listed in Table S3 (*SLC6 A16*) had a statistically significant increase in the number of on-ART IS relative to the PBMC IS, suggesting that there might have been selection for cells with proviruses in the gene, despite their opposite-sense orientation, although there was no obvious clustering of the on-ART IS relative to the PBMC IS (Figure S3 C, D).

We also considered the possibility that there could be regions outside of genes where proviruses could provide a selective advantage to the infected cell. We used an unbiased scan to look for evidence of clustering of proviruses within 10-kb regions in the on-ART dataset (Figure S4, Table S4). With the exception of genes that had already been described, all of the clusters that were found in the on-ART dataset were also present in the PBMC dataset. Sites in which there was clustering in both the in vitro and in vivo dataset are regions that are favored for integration “hot spots” – this is in contrast to the clusters that arise in vivo from post-integration selections. Not surprisingly, clusters of proviruses were found in some of the genes that are the most favored targets for integration, including *PACS1* (Figure S4F), the gene which had the largest number of sites in the infected PBMC (Figure 2A, Inset), and the lncRNA genes *NEAT1* and *MALAT1* (Figure S4E), which are among the genes with the highest density of IS in vivo and in vitro (Table 2). One of the hot spot clusters identified in both the PBMC and on-ART datasets appeared to be intergenic (Figure S4, panel H). However, analysis of the RNA-seq data showed that there is a transcription unit in this region which has not yet been annotated.

### Clonal amplification of HIV-infected cells

In most of the analyses presented thus far, we collapsed all the breakpoints associated with each IS to one, a step that simplified the comparisons of the in vitro and in vivo data. One of the most striking features of the distribution of HIV IS in vivo is the presence of identical IS that derive from clonally-amplified cells. The effect of clonal amplification on the IS data can be seen by comparing the distribution of IS in the MKL2 cluster in Figure 4B before (bottom panel) and after (middle panel) the amplified sites were collapsed down to unique sites. In this case, the 30 unique IS were found a total of 144 different times – an average amplification ratio of nearly 5, ranging from 1 (no amplification) to more than 25. Overall, the amplification ratio for all sites across the entire genome was 1.6 for the on-ART samples, and 1.2 for the pre-ART samples, confirming that infected cells can grow into clones before ART is initiated [18] (Figure 6). The amplification ratios were independent of whether the proviruses were inside or outside of a gene, the orientation of the proviruses relative to the host genes, and of the level of expression of the host gene (Figure 6 and S5). The amplification ratios for the proviruses in the six genes where proviruses could be selected, although quite variable, were, on average, not significantly different from all the other genes (Figure 5B). When the amplification ratios for each site were ranked and plotted in decreasing order, the sizes of the clones in the six genes were scattered across the distribution (Figure S6). Thus, the degree of clonal amplification cannot be taken as evidence of positive selection for a provirus in a specific location, and such proviruses are not a significant contributor to the overall level of clonal expansion.

**Figure 6.**
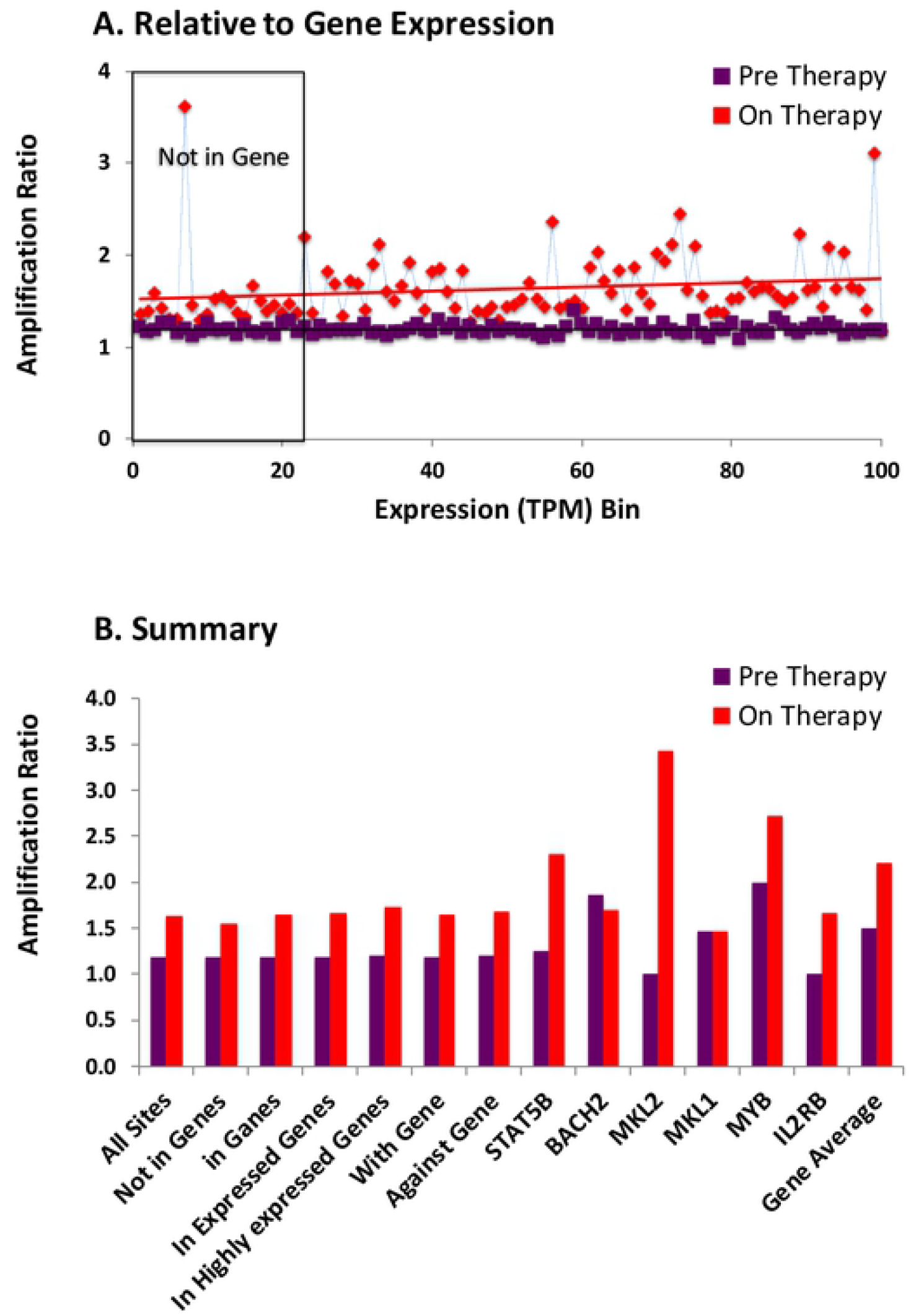
IS distribution, RNA level, and clonal amplification. The amplification ratio for all integrated proviruses was calculated using (total number of IS)/(number of unique IS). **A.** Each RefSeq gene in the non-overlapping dataset was assigned to one of 100 bins (ca. 200 genes in each bin) based on the level of RNA for the gene (as TPM in the in vitro infected PBMC), and the overall amplification ratio was calculated for the sites in each bin. Non-expressed genes were assigned to bins at random. The regression lines shown have a slope of 0.0008 and 0.0013 (p=0.88 an 0.12) for pre-ART and on-ART bins, respectively. **B** shows the clonal amplification ratios pre-ART and on-ART. Color coding is the same as in previous figures: Plum, pre-ART; red, on-ART.

### Integration in centromeric repeats

The complex nature of the alphoid repeat sequences precludes precise mappting of IS in centromeres, and, for this reason, they are not annotated in the hg 19 (or hg38) genome database and were not detected using our the standard pipeline to identify and locate IS, even though the cetromeres comprise approximately 7% of the genome [40]. [See, for an example, the large gene-free area around 40 Mb of chromosome 16 (Figure 3)]. It has recently been reported that in rare HIV-infected individuals who maintain very low levels of viremia in the absence of ART (known as elite controllers) the intact proviruses are preferentially found within centromeric sequences [41]. We used a manual protocol to search specifically for centromeric IS in a portion of the data from the PBMC and on-ART datasets. Although indentifying IS in centromeres is not trivial, and we are not confident that we identified all the centromeric IS in the datasets, Table 4 shows that there are detectable IS in the centromeric repeats in both the PBMC and on-ART datasets. The data show that the amplification ratio of the on-ART proviruses (1.70) is not significantly different from the proviruses found outside genes (1.62, P=0.77). Nor is the amplification ratio in centromeric proviruses different from proviruses that are integrated in genes (Figure 6). Proviruses integrated in centromeric regions do not appear to be expressed and there are no known genes in centromeres. Thus, these results support our conclusion that factors other than the effects that proviruses have on the expression of genes in which they are integrated are the primary factors that control the clonal expansion of infected cells.

**Table 4.** IS detected in centromeric repeat DNA^a^.

|  | Genome<br>(Mb) | PBMC<sup>c</sup><br>unique<br>sites | Sites on therapy<sup>d</sup> |  |  |
| --- | --- | --- | --- | --- | --- |
|  |  |  | Total | Unique | Amp<br>ratio |
| All | 3,080 | 216,054 | 64,283 | 37,967 | 1.69 |
| Outside genes<sup>b</sup><br>(%) | 524<br>(17%) | 36,729<br>(17%) | 14,142<br>(22%) | 8,732<br>(23%) | 1.62 |
| In centromeres<br>(%) | 216<br>(7%) | 178<br>(0.082%) | 129<br>(0.20%) | 76<br>(0.20%) | 1.70 |

<sup>a</sup> Data from additional assays of some samples (Table S1) were analyzed for IS in centromeric repeat DNA.
<sup>b</sup> Recalculated using the proportions in Table 1 to estimate the numbers of total and unique IS for the different-sized samples.
<sup>c</sup> A portion of the data from PBMC infected in vitro. Because “amplification” is due only to DNA replication immediately following infection, it is without significance here and is not reported.
<sup>d</sup> From the donors (Table S1), including additional samples that were taken after those reported in the rest of the manuscript.

## Discussion

HIV persists for decades on fully suppressive ART, primarily as poorly expressed proviruses in infected cells that are long-lived, and continue to divide. Many (perhaps all) of these long-lived cells are in expanded clones. The present study was undertaken to better understand the factors that affect which provirus-containing cells survive in vivo. We compared a large IS dataset obtained from PBMC infected in vitro to pre-ART and on-ART datasets compiled from a number of in-house studies of HIV-infected individuals. The approach we used [21, 30, 31] provided IS data that are unbiased by the distribution of restriction enzyme sites in the human genome. The data were extensively screened to remove cross-contamination and any PCR artifacts or other errors.

There are three factors that affect intial distribution of proviruses in vivo: 1) negative selection against cells with proviruses in highly-expressed genes; 2) positive selection for cells with proviruses in specific regions of certain genes; and 3) stochastic effects, probably due to immune signaling, that lead to clonal amplification of some infected cells. The primary determinant of the pre- and on-ART distribution of IS is their initial distribution at the time the cells were infected. In the experiments we report here, that distribution was modeled using the distribution of IS in donor PBMC infected in vitro. In general, the three IS datasets are very similar. The excellent agreement, for most genes, in the distribution of the IS in the PBMC data and the two in vivo datasets supports our use of the PBMC IS dataset to accurately reflect the initial distribution of IS in vivo. The pre-ART IS distribution more closely matches the PBMC IS distribution than the on-ART dataset because the pre-ART dataset is a mixture of newly infected cells and cells that have been infected for longer times. The ways in which the PMBC and on-ART IS datasets don’t match provide insights into the selective forces that reshape the intial distribution of IS in vivo.

As has been known for many years, HIV DNA integration strongly favors the bodies of transcribed genes and the orientation of the provirus is independent of the chromosomal orientation of the host gene [7-9]. The strong association of HIV integration with expressed genes is shown in Figures 1 and 2 and Table 1, where the normalized integration density (IS/Mb) in expressed genes was more than 15-fold greater than in the rest of the genome. Integration density in PBMC was 8-fold greater in genes with high expression levels than in those expressed at low levels.

Many highly-expressed genes are good targets for HIV integration; however, high levels of gene expression do not always predict high levels of integration. This effect may be mediated by the distribution of LEDGF on chromatin [9, 42-44]. For example, in Figure S2, which shows three adjacent *STAT* genes, all of which produce similar levels of RNA in PBMC. *STAT5B* is a good target for HIV integration, *STAT3*, less so, and *STAT5A* has very few IS. These discrepancies may reflect the presence, in the CD8-depleted PBMCs used for the RNA-seq analysis, of differentially expressed genes in cells that are and are not targets for HIV infection, or genes that express very stable transcripts, so that the steady state RNA levels measured by RNA-seq are not indicative of the level of transcription, and there could be be other explanations.

Almost all regions of the host genome that are not expressed are poor targets for HIV integration. This result was consistently seen in both in vitro and in vivo. An unbiased search for clustered IS turned up one apparent intergenic region, on chromosome 6, in which there was a cluster of IS but no annotated gene. However, the RNA-seq data show that this region is transcribed, although not yet annotated as a gene. This result suggests that clusters of HIV IS might be a useful way to check genome databases for genes that have not been annotated. Very rarely, we found annotated genes that were good targets for integration both in vivo and in vitro, in the absence of detectable levels of RNA in PBMC. *HORMAD2*, whose protein product is involved in meiosis, is a good example (Figure S5), in that it had the largest number of proviral ISs of any non-expressed gene in both the in vitro-infected PBMC and on-ART datasets. Interestingly, but perhaps coincidentally, a provirus in this gene (in the form of a solo LTR) is in one of the largest clones we have seen in any on-ART donor [21, 45].

Although the distributions of IS in genes in the PBMC, pre-ART, and on-ART datasets are very similar, some clear differences emerged when the frequency of the IS was plotted against the level of gene expression (Figure 1). The normalized fractions of IS in the three datasets were identical for genes that were poorly expressed; however, the fraction of the IS in highly-expressed genes was higher in the in vitro PBMC dataset than in either the pre-ART or the on-ART datasets. The in vivo datasets also showed that there is selection against proviruses in either orientation in highly-expressed genes. Although an overall against-the-gene orientation bias has been previously reported for on-ART samples [21, 29], the fact that the magnitude of the selective effect depends on the level of expression of the genes had not been reported. Nor was it previously known that there is a weaker selection against proviruses in highly-expressed genes that are oriented opposite to the gene.

It has been proposed that latent, intact, proviruses within genes may be more likely to be expressed than those in extragenic regions, leading to death of the cell and accounting for a bias for defectiveness for proviruses in genes [24], although the net effect is not large and counterexamples have been reported [46]. Although the vast majority of proviruses are defective, that type of selection could also apply to cells with defective proviruses if the defective proviruses expressed epitopes that are reconized by CTL, for which there is conflicting evidence [47-49] Given the recent results showing that only a small fraction of cells, containing either intact or defective proviruses, express unspliced viral RNA [25], we think it more likely that the selection against cells with proviruses in highly-expressed genes is due to the effects of the provirus on the expression of the host gene. In the case of proviruses in the same orientation as the gene, these effects would involve the insertion of viral sequences containing signals for transcription initiation, splicing, polyadenylation, etc., interfering with its expression. Certain endogenous proviruses of mice are also known to affect the expression of the host gene in which they reside [50, 51]. The effect on gene expression is limited to the allele in which the provirus is inserted, possibly explaining why the strongest selection is seen for proviruses that are integrated in the most highly-expressed genes. Proviruses integrated in the opposite orientation to a gene could also reduce its expression by transcriptional interference [52]; however, the effects on gene expression are likely to be weaker for proviruses in the opposite orientation to the gene. In support of the interpretation that the effect is on the level of expression of the host gene, the effect is due to a weak selection on a large number of genes, rather than strong selection in a small number of genes. The observation that there is a much weaker relationship between gene expression and provirus loss in the pre-ART than in the on-ART samples (compare Figure 1, panels C and D) also supports this conclusion.

It has been known for nearly 40 years that modification of gene structure and expression by retroviral DNA integration can cause cancer in animals [53, 54]. Provirus-mediated effects are known to induce oncogenic modification of genes by the insertion of a promoter or an enhancer, which leads to the overexpression of a host oncogene and/or a truncation of the encoded protein [55]. Similar mechanisms likely affect the expression of the three genes *(BACH2, MKL2,and STAT5B)* previously identified as targets for HIV provirus mediated preferential expansion or survival of infected T cells in humans [21, 23, 28, 37, 56]. In the present study, we did a comprehensive search for additional genes in which an HIV provirus insertion could cause similar posititve selection for infected T cells. Host genes in which a provirus could help the cell proliferate or survive were identified by comparing the IS frequencies in individual genes in the on-ART dataset and the PBMC dataset. Genes were identified based on finding a large increase in the integration frequency in vivo. In the genes in which there was selection for proviruses, there was also strong selection for proviruses in the same orientation as the gene, and the proviruses were clustered in one or a few introns.

Using the same criteria, we found three additional genes in which proviruses contibuted to the growth and survival of the cell: *MKL1, IL2RB*, and *MYB*. Because we began with a large on-ART dataset, this set of six represents a complete, or nearly complete, set of the host genes in which enhancement of growth and/or survival of the host cell by an integrated HIV provirus is likely to play an important role in HIV persitence. As described in Results, there were only 10 (out of 33,000) unique ISs in *MYB* in the on-ART dataset. While it is possible that there are other poorly-targeted genes in which HIV proviral integration would have similar effects, any gene that has not yet been identified must be a very poor target for HIV integration. Of the 20,207 genes in our modified database, only about half (11,422) had one or more IS in the PBMC dataset, and only about one quarter (5159) had more than 10. Any gene we might have missed would have been even rarer than *MYB*, and would have made only a negligable addition to the list of genes in which proviruses can make a contribution to the growth or survival of the host cell.

Proviruses integrated in one of these six genes comprised, together, approximately 1.2% of the total unique proviruses. Many of the proviruses in these genes are defective (Hill, S. and Maldarelli, F, unpublished observations); there are no published reports of an intact provirus in any of these six genes. If clonally expanded cells are counted, the cells with proviruses in the six genes comprise about 1.5% of the total infected cells. Thus, it is unlikely that the cells that have expanded because they have a provirus that drives their proliferation and/or survival are an important part of the viral reservoir, which, by definition, is composed of intact proviruses.

We and others have previously pointed out that a large fraction of the HIV proviruses in on-ART samples are integrated in genes/oncogenes that play important roles in the proliferation and survival of cells [21, 23, 57]. However, as the data we present here clearly show, this preference is a result of the propensity of HIV proviruses to integrate in expressed genes, and not due to post-integration selection for cells that have proviruses integrated in oncogenes other than the six that have already been discussed. For all other oncogenes, the fraction of the IS in the gene does not increase when the PBMC data and the on-ART data are compared, which shows that there is not a strong post-integration selection in vivo for proviruses in other oncogenes.

As previously mentioned, proviruses of oncogenic retroviruses can modify the expression of host genes [54, 55], and it is likely that HIV proviruses can use similar mechanisms to alter host gene expression. In *BACH2, STAT5B*, and *IL2RB*, the selected proviruses are in introns upstream of the start site for translation, and probably provide a new promoter for the gene, and a new upstream leader sequence that is derived from the provirus [28]. Proviruses integrated in *MKL1* and *MKL2* interrupt the protein coding region, which if the mechanism of activation is promoter insertion, would lead to the expression of truncated proteins that lack amino terminal domains present in the normal proteins. In the case of *MYB*, the proviruses are in a large 3’ intron and are likely to cause premature polyadenylation, resulting in a protein product that lacks the correct 40 C-terminal amino acids. The inserted proviruses would also lead to the synthesis of an mRNA lacking the long 3’ noncoding region of *MYB*, which has been reported to contain signals that promote rapid RNA degradation [58]. These are all mechanisms that oncogenic retroviruses use to modify expression of host genes [54].

The cells in which an HIV provirus has integrated into one of the six oncogenes have a growth and/or survival advantage, but they are not cancer cells. However, as with cancer, provirally induced misexpression of the host target gene product may promote cell division. Alternatively, the altered protein may provide a survival advantage, by, for example, suppressing apoptotic signaling, which is a normal part of the T cell immune response. Of the six genes in which proviruses led to positive selection, only three were in cells that were clonally amplified to a greater extent than average for all cells in the on-ART samples (Figure 5C). It is possible that this difference reflects two different types of positive selection: Cells in which the provirus is in *STAT5B, MKL2*, and *MYB* may have increased potential for growth and cells with proviruses in *BACH2, MKL1*, and *IL2RB* may have better long term survival due to a reduction in apoptotic signaling.

Clonal amplification is the most striking feature of persistent proviruses on-ART, and is responsible for the majority – perhaps all – of the infected cells that persist on long-term ART, whether they contain intact or defective proviruses. We show here that provirus-driven clonal expansion account only for a small minority of the clonally expanded cells, which means that forces independent of the location and structure of the provirus cause the majority of the clonal expansion that characterizes infected cells in indivduals on ART. Thus, in most cases, the proviruses only provide useful markers for the cells in expanded clones. In support of this conclusion, we found that the extent of clonal expansion is independent of the location of the provirus, whether it is in a gene or intergenic, the expression level of the host gene, and the orientation of the provirus relative to the gene (Figure 5). Even when integration is in centromere-associated alphoid repeat DNA, which is believed to be highly inimical to proviral expression [41, 59], the extent of amplification was not different from what was found for all of the proviruses taken together. We therefore conclude that proviral-driven expression of “cancer genes” plays only a minor role in the clonal expansion of infected cells, and that other mechanisms, probably antigen-driven or homeostatic expansion, account for the proliferation and persistence of the large majority of clones of infected cell in indivduals on ART, especially those with intact proviruses.

The differences in the distributions of the proviruses in the PBMC infected in vitro and in the pre- and on-ART samples, although highly significant, are small (Figure 2). This result implies that the location of the proviruses in the host genome has only a small effect on the survival of an infected cell. Because cells that carry expressed proviruses are preferentially lost on ART, we conclude that the location of a provirus in the host genome has little effect on the expression of the provirus in long-lived cells. We show that there is a modest loss of cells with proviruses in highly expressed genes, and the loss is greater for the provirus in the same transcriptional orientation as the gene. However, as we have already discussed, this selection is more likely to be due to the effects of the proviruses on the expression of the host genes than the other way around. Turning the problem around, if the cells that carry proviruses that are not expressed have a survival advantage, and if the location of the proviruses in the genome was a substantial factor in whether the proviruses were expressed, we would have expected to see a larger shift in the distribution of the IS on ART. Although there is, with time on ART, a very slow and erratic selective loss of relatively intact or inducible proviruses [47, 60, 61] which leads to an enrichment in the fraction of defective proviruses [45], there is no evidence that the location of the provirus plays a role in this selection. Our conclusion that the location of an HIV provirus in the genome has little, if any, effect on its expression in infected individuals is supported by data, obtained from cells that were infected with HIV-based vectors in vitro, that the distributions, in the host genome, of the vector proviruses that are and are not expressed are quite similar [59].

In conclusion, the distribution of HIV proviruses, even after years of complete virological suppression, in which the majority of the infected surviving cells are removed from the originally infected cells by many divisions, still closely resembles the distribution of the proviruses at the time the cells were initially infected, represented by the proviral distribution in PBMC acutely infected in vitro. In addition to a modest modification to the initial distribution by a positive selection for proviruses integrated in the six genes we have listed, the distribution is also modified by selection against cells with proviruses integrated in highly-expressed genes. There is also extensive clonal amplification; however, more than 98% of the amplification results from factors in which the proviruses and their locations in the genome play no apparent role in the behavior, growth, or survival of the infected cells.

## Methods

### Donors and cells

PBMC were isolated from 120 mL of blood from two anonymous healthy, HIV negative donors and depleted of CD8+ T cells using Dynabeads (ThermoFisher). Approximately 30 x 10^6^ cells were placed in 30ml of R10 medium, were treated with 0.5ug/mL of phytohemagglutinin (PHA) [62], and were incubated at 37°C for 2 days. Both donors provided written informed consent and the donation protocol was approved by the University of Pittsburgh Institutional Review Board.

Aliquots of 10^7^ cells, treated with PHA (0.5 μg/ml) were pelleted and resuspended in 2mL of fresh RPMI-1640 media supplemented with 10% FBS and containing 10^7^ IU/ml HIV-1_BAL_ [63] for a multiplicity of infection of 1 infectious unit/cell as assayed in GHOST cells [64]. After 2 hours, the cells were washed and incubated at 37°C for a further 2 days. 10^7^ cells from each flask were pelleted, resuspended in 1ml fresh cryopreservation medium, and stored at −80°C.

PBMC were isolated from blood drawn from HIV-infected donors participating in one of a number of clinical studies under the aegis of the HIV DRP, the University of Pittsburgh, and Stellenbosch University as described in the references to Table S1. All donors (or their guardians) provided written informed consent, and all studies were approved by the local institutional review board.

### Integration site analysis

The distribution of HIV IS in all PBMC samples was determined by linker-mediated nested PCR of fragmented DNA using LTR and linker-specific primers and paired-end Illumina sequencing as described [21]. Raw data were analyzed using the bioinformatic pipeline described by Wells et al [31], which includes steps designed to remove artifacts due to mispriming, cross contamination, etc. such as those found in Cohn et al. [31, 36]. When we compared the global distribution of the PBMC IS from the two donors, the correlation coefficient (*r*) was 0.98 and, for all analyses, we combined the data from the two donors, giving about 385,000 total IS. All IS data have been posted on the Retroviral Integration Database (RID) at https://rid.ncifcrf.gov/ [65], and can be found using the PubMed ID of this paper. All data are also present in the Microsoft Excel document that can be found as Supplemental File 1. The PBMC dataset was also used as a comparator for a study on SIV IS distribution in the macaque model [29].

The IS analysis pipeline used for this study reports the site (at the end of the 3’ LTR) in the hg19 human sequence database, the orientation of each provirus relative to the host chromosome, and the number of differently sheared flanking host DNA sequences (or breakpoints) associated with each distinct IS. The breakpoint count indicates the relative number of clonally-expanded cells containing the same provirus. For many analyses, the clonally expanded IS data were collapsed so that only the unique IS were used. To map IS to centromeric regions, fastq files generated from the Illumina platform were processed and analyzed using our in-house standard pipelines for HIV integration sites analysis, with specific modifications to accommodate centromere mapping. Reads properly trimmed and filtered were subjected to centromere alignment using BLAT without -ooc filter to GRChg38 (released Dec. 2013) centromere assembly obtained from the University of California Santa Cruz genome browser (https://genome.ucsc.edu) as a reference. Reads mapped to centromeres were further merged and filtered to identify real integration sites. Manual inspections were used to finalize the mapping results.

Gene locations were obtained from the RefSeq set [35]. For counting purposes, in cases of overlapping genes, the 5’-most gene was truncated and genes entirely inside other genes were removed so that all sequences in host genes were assigned to a single gene. This correction was necessary to avoid double counting of IS where there are overlapping genes.

The maps shown in Figures 3-5 and S3-S5, were generated using a custom-made Microsoft Excel workbook, as described in Figure S2, which also contains all the data used in this paper, and allows the user to select up to 5 datasets at a time, and choose a specific gene, chromosome, or chromosomal region to be viewed. The program is available as Supplemental File 1.

P values for most comparisons were calculated using a binomial distribution or Fisher’s exact test, as appropriate. P values for the comparisons between selected genes and total IS in Figure 6 were done using a bootstrap analysis (10^6^ replicates each).

### RNA-seq analysis

Total nucleic acid was extracted from CD8-depleted donor PBMC infected with HIV following PHA stimulation. RNAseq was carried out using an Illumina TruSeq mRNA library kit with 200ng of poly(A) selected RNA as input. Libraries were pooled and sequenced on Illumina NextSeq500 with TruSeq V2 chemistry paired end reads x 75bp, yielding ~100 million reads per sample. Sequencing was performed at NCI CCR Sequencing Core Facility. RNAseq data analysis was performed using the standard RNAseq workflow in CLC Genomics Workbench V12. Reads were mapped to human genome hg19 and Ensembl genes annotation v74 was used to calculate gene expression values as transcripts per million (TPM). Only protein coding genes and lncRNA genes were included for combined gene expression and IS analysis.

## Acknowledgments

The authors would like to thank Teresa Burdette for help in preparing the manuscript, Allen Kane for help with the figures, and Anne Arthur for editorial help. This research was supported by the Intramural Research Programs of the NIH, National Cancer Institute. JMC was a Research Professor of the American Cancer Society and was supported by the NCI through a Leidos subcontract, l3XS110 and research grant R35 CA200421. JWM was supported by the NCI through a Leidos subcontract, 12XS547. This project has been funded in whole or in part with federal funds from the National Cancer Institute, National Institutes of Health, under Contract No. HHSN261200800001E. The content of this publication does not necessarily reflect the views or policies of the Department of Health and Human Services or the National Institutes of Health, nor does mention of trade names, commercial products, or organizations imply endorsement by the U.S. government.

## Data and materials availability

All data used for this study are contained in Supplemental File 1, along with the Microsoft Excel based mapping tool used to generate Figures 3-5 and S2-S5. The IS data have also been deposited in the Retroviral Integration Database (https://rid.ncifcrf.gov/), and can be accessed using the ID for this paper.

## Supplemental Materials

**Figure S1.** Gene expression and integration.

**Figure S2.** Application for visualization of IS and expression data.

**Figure S3.** Genes with high frequency of proviruses integrated in the opposite **transcriptional** orientation.

**Figure S4.** Clusters of IS not associated with cell growth or survival.

**Figure S5.** Integration into a non-expressed gene.

**Figure S6.** IS Ranked by Clonal Amplification.

**Table S1. Sources of Donor IS Data.**

**Table S2. Distribution of IS as a function of gene expression (TPM)**

**Table S3. Genes in which proviruses oriented against the host gene were strongly selected in the on-ART dataset**

**Table S4**. **Genes with the largest clusters in 10 kb windows in on-ART samples**

**Table S5 IS detected in centromeric repeat DNA**

**File S1. Integration-Transcription Maps Data and Worksheet**

